# Evolution in salmon life-history induced by direct and indirect effects of fishing

**DOI:** 10.1101/2021.01.08.425869

**Authors:** Y. Czorlich, T. Aykanat, J. Erkinaro, P. Orell, CR. Primmer

## Abstract

Understanding the drivers of evolution is a fundamental aim in biology. However, identifying the evolutionary impacts of human activities, both direct and indirect, is challenging because of lack of temporal data and limited knowledge of the genetic basis of most traits^1^. Atlantic salmon is a species exposed to intense anthropogenic pressures during its anadromous life cycle^2^. Previous research has shown that salmon age at maturity has evolved towards earlier maturation over the last 40 years, with an 18% decrease^3^ in the allele associated with late maturation at the large-effect *vgll3* locus^4^; but the drivers of this change remain unknown. Here, we link genetic and phenotypic changes in a large Atlantic salmon population with salmon prey species biomass in the Barents Sea, temperature, and fishing effort in order to identify drivers of age at maturity evolution. We show that age at maturity evolution is associated with two different types of fisheries induced evolution acting in opposing directions: an indirect effect linked with commercial harvest of a salmon prey species (capelin) at sea (selection against late maturation), and a direct effect due to temporal changes in net fishing pressure in the river (surprisingly, selection against early maturation). Although the potential for direct and indirect evolutionary effects of fishing have been acknowledged, empirical evidence for induced changes at the genetic level has been lacking^5^. As capelin are primarily harvested to produce fish meal and oil for aquaculture^6^, we hereby identify an indirect path by which Atlantic salmon aquaculture may negatively affect wild populations.

## Main text

Identifying biotic and abiotic factors linked with phenotypic change is common but cases demonstrating evolutionary (i.e. genetic) effects remain rare ^1,7^. Changes due to indirect effects, whereby the impact of one species on another is mediated by a third species, are prevalent in nature and can have important ecological and evolutionary consequences ^8^. Indirect effects are expected to be of more central importance in current-day ecosystem dynamics due to the growing anthropogenic pressure and climate change rate which can strongly influence species, and interactions between them, in complex ways e.g. ^9^. However, detecting indirect evolutionary impacts is extremely challenging in natural ecosystems and has often been neglected^8^.

An example of human activity that can have profound effects, both directly on the target species, and indirectly on entire ecosystems, is fish harvest ^10^. Fishing is generally performed at high intensity, over prolonged periods of time, and can be trait selective. It can consequently induce evolution of traits such as size and age at maturity in the target species ^5,11^ but also induce larger, ecosystem level, changes, e.g. by reducing the abundance of predators, prey and/or competitors ^12^. To date, most research has focused on the effects of harvest on target species at the phenotypic level and cases demonstrating direct effects at the genetic level are rare, as are empirical examples of indirect evolutionary impacts ^5,13^. Knowledge of indirect ecological and evolutionary effects is critical for properly evaluating the consequences of different fisheries management strategies ^10^.

Atlantic salmon (*Salmo salar*) have a complex life-history, utilizing both freshwater and marine habitats and thus affect, and are affected by, multiple ecosystems ^2^. Sea-age at maturity in Atlantic salmon (the number of years an individual spends in the marine environment before returning to fresh water to spawn), or sea age, is an important life history trait with an evolutionary trade-off between survival and reproduction. Later maturing individuals are larger and have higher reproductive success ^14^ but run higher risk of mortality before spawning. Age at maturity has been associated with a major effect locus located in the genome region including the *vgll3* gene that explains 40% of the variation in the trait ^4^. We recently demonstrated adaptive evolution towards younger age at maturity at the *vgll3* locus over 40 years in a large salmon population in northern Europe ^3^. Both sexes experienced a decline in the *vgll3* allele linked with later maturation and only males showed a clear evolutionary response at the phenotypic level, due to *vgll3* sex-specific allelic and dominance effects ^3,4^. However, the environmental drivers of the evolution in age at maturity remained unknown.

Here we identify environmental drivers of phenotypic and adaptive changes in age at maturity in the same Atlantic salmon population from the Teno river, in far north Finland and Norway ^3^. We investigated ecological and environmental variables potentially affecting the relative fitness (a combination of marine survival and reproductive success) of salmon with different maturation ages, and therefore sizes: fishing effort at sea, fishing effort in the river, sea temperature, and abundance of three key prey species for salmon during their marine migration in the Barents sea ecosystem (capelin *Mallotus villosus*, herring *Clupea harengus* and krill, Extended Data Fig. 1). We used (quasi)binomial generalized linear models (GLM) to identify environmental variables linked with temporal variation in *vgll3* allele frequencies, and thereby age at maturity, in a 40 year time series (1975 – 2014) consisting of 1319 individuals assigned to the Tenojoki population in ^3^ (see methods, Supplementary notes). These analyses indicated that annual riverine net fishing license number (used here as a proxy for annual fishing pressure, see Methods) had the strongest effect on *vgll3* allele frequency (Fig. 1). Surprisingly, annual riverine fishing pressure was positively associated with the *vgll3*\**L* allele frequency (standardized regression coefficient 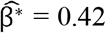, F_(1)_ = 27.79 and P-value < 0.001) indicating that higher net fishing pressure in the river resulted in higher frequencies of the allele associated with later maturation in salmon and therefore larger size (Extended Data Table 1). We further verified the results using a residual regression approach (de-trending) to account for potentially confounding factors creating temporal trends in the dependent and independent variables, and thus avoid possible spurious associations ^15^. The association between annual fishing license number and *vgll3* remained in the de-trended model including a significant, negative, year effect e.g. ^3^, indicating that fishing pressure is also linked with annual *vgll3* allele frequency changes around the trend (F_(1)_ = 14.15 and P-value < 0.001, Fig. 1, Extended Data Table 1). Capelin biomass in the Barents Sea was also positively associated with the frequency of the *vgll3* allele associated with later maturation and larger size (*vgll3*L)* in salmon in both the normal (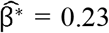, F_(1)_ = 20.77 and P-value < 0.001) and de-trended models (F_(1)_ = 10.64 and P-value = 0.001, Extended Data Table 1). Herring and krill biomass also had a significant effect on *vgll3* allele frequencies in a similar direction to capelin (Fig. 1, Extended Data Table 1), however, they did not remain significant in the de-trended model. There was little evidence for associations between the other variables and *vgll3* in the GLM (Extended Data Table 1).

**Figure 1:**
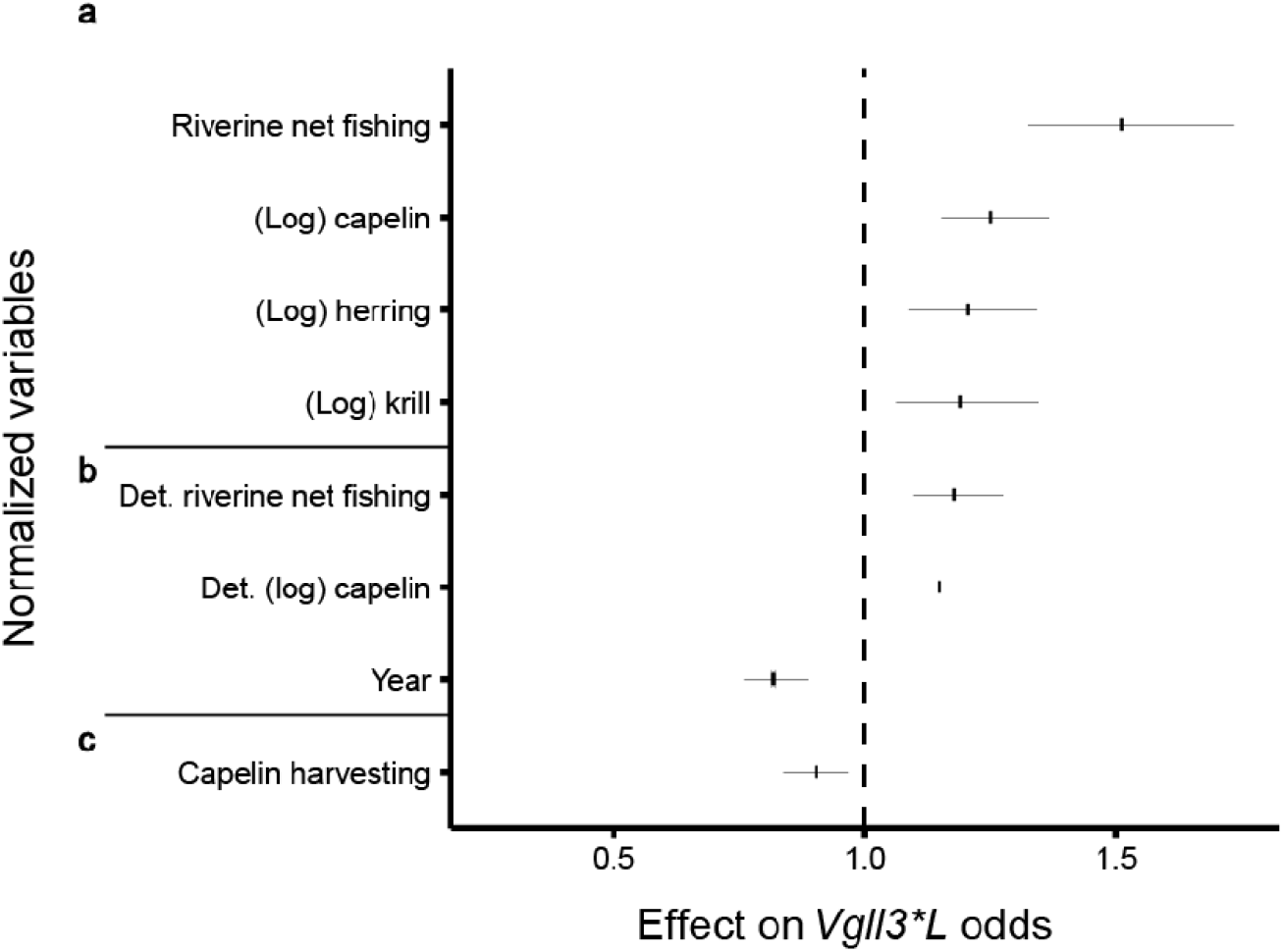
Standardized regression coefficients for variables significantly associated with *vgll3*L* odds (later, larger maturation). The estimates come from the ***a***, initial quasi-binomial model. ***b***, de-trended model. ***c***, Monte Carlo Method for assessing mediation (i.e. indirect effects). The dotted line indicates no effect on *vgll3*L* odds. The error bars correspond to 95% confidence intervals.

Analyses at the phenotypic level also supported the influence of prey biomass and fishing on Tenojoki Atlantic salmon fitness. After controlling for the *vgll3* sex-specific genetic effects, a multinomial model indicated that the probability of observing older, later maturing Atlantic salmon in the river increased with capelin, herring and krill biomass (Supplementary notes, Extended Data Table 3). This result is expected if the abundance of these prey species is positively associated with salmon survival at sea, as only survivors returning back to the river are sampled. However, it can also reflect plastic changes of maturation probabilities. This model also showed that a higher number of riverine net fishing licenses was associated with a higher probability to observe late maturing salmon at the end of the fishing season (Extended Data Table 3), which is expected if net fishing targeted preferentially small, early maturing salmon carrying *vgll3*\**E*. Given that the sea-age at maturity of Tenojoki females is, on average, considerably older than males, and therefore the time spent in the marine environment longer (2.8 vs 1.5 years), environmental conditions strongly affecting marine survival are also expected to influence the sex-ratio of adults returning from their migration to spawn. In accordance with this prediction, a binomial model showed that the proportion of returning females increased with prey biomass and riverine net fishing (Supplementary notes; Extended Data Fig. 2, Extended Data Table 3). The effects of net fishing on sex-ratio and age at maturity probability, however, were not significant anymore in the detrended regressions (Extended Data Table 2, Extended Data Table 3).

Forage fishes like capelin have important roles in marine ecosystems by enabling energy transfer between lower (plankton) and upper (predators such as large fish, seabirds and mammals) trophic levels ^6^. In the Barents Sea, the capelin stock experienced several dramatic collapses during the 40-year study period (Extended Data Fig. 3) due to overexploitation in commercial fisheries combined with predation by herring and cod ^16^. We therefore quantified the potential indirect effects of capelin harvesting (while accounting for other ecosystem interactions) on *vgll3* allele frequency dynamics using a multispecies Gompertz model developed in ^17^ (Extended Data Fig. 1, Extended Data Fig. 3, Extended Data Fig. 4). The analysis indicated a significant indirect effect of capelin harvest rate on age at maturity evolution in Tenojoki salmon, with a 30% decrease in the *vgll3*\**L* allele odds per harvest rate unit (Monte Carlo method for assessing mediation: CI_95%_ = [0.116, 0.471]). The evolutionary effect of capelin harvest was the strongest during the early years of the time series, likely because of repeated capelin fisheries closures and fishing effort reduction later on, following the capelin stock collapses (Fig. 2; Extended Data Fig. 5a). This is the same time period when most of the evolutionary changes in male age at maturity occurred ^3^.

**Figure 2:**
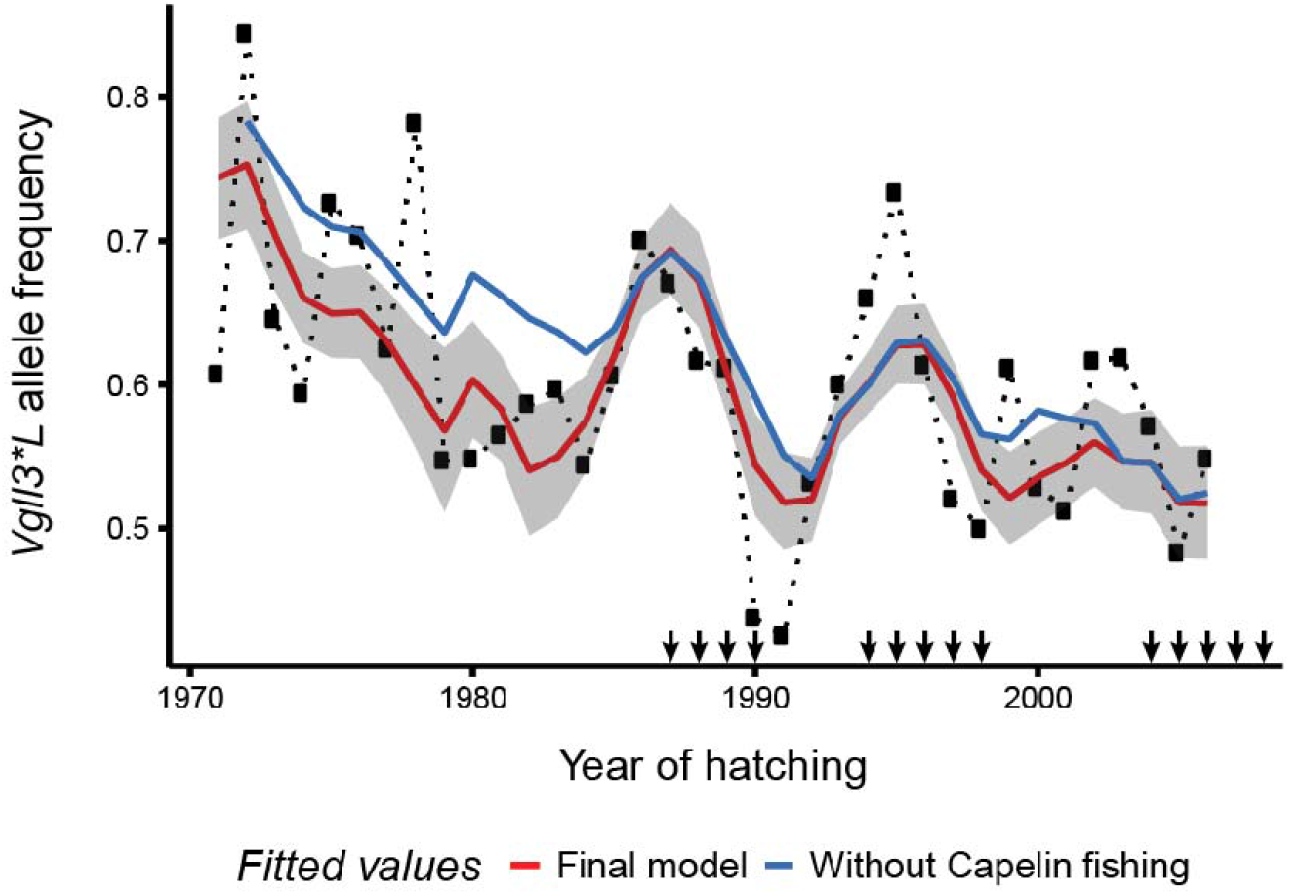
Temporal changes in *vgll3*L* (later maturation) allele frequency. The black dotted line represents the observed data, the red line the allele frequency averaged from model individual fitted values and the blue line the expected *vgll3*L* allele frequency assuming no capelin fishing (annual effect on returning salmon). Black arrows indicate years without capelin harvesting (harvest rate < 0.5 percent). Shaded areas indicate 95% bootstrap intervals based on 3000 replicates (see methods for details).

The direction of the selection pressure in the riverine net fishery may seem counter-intuitive as generally, net fishing is thought to select against large individuals e.g. ^18,19^. However, net fishing in Tenojoki includes the use of multiple net types with different selectivity: weir, gillnet and driftnet^20^. Sonar data enabled an assessment of the selectivity of each net type (see Methods), and indicated that weir fishing selects against the early maturation allele (*vgll3*E*) by capturing proportionally more smaller, and earlier maturing, individuals whereas driftnet and gillnet select against the *vgll3*L* allele (Extended Data Fig. 6, Extended Data Fig. 7). The positive effect of riverine net license number on the *vgll3*L* allele frequency may thus be explained by the predominant use of weirs (representing 54% of net catches on average). Over time, the net fishing selective pressure against the *E* allele is expected to have decreased through a dramatic reduction in the net fishing effort during the time series due to stricter fishing regulations (e.g. net fishing licences and number of weirs were >30% lower in 2014 vs. 1976, Extended Data Fig. 5b) and in selectivity due to changes in the relative use of the fishing gears, size distribution and sex-ratio (Fig. 3, Supplementary notes). Overall, the extra mortality at sea of late maturing *vgll3*L* individuals, which varied over time according to prey biomass, was not sufficiently compensated by their (decreasing) size-induced survival advantage due to riverine net fishing, nor their reproductive advantage due to larger size ^14^, thus resulting in the observed overall decrease in the *vgll3*L* allele frequency (Fig. 2).

**Figure 3:**
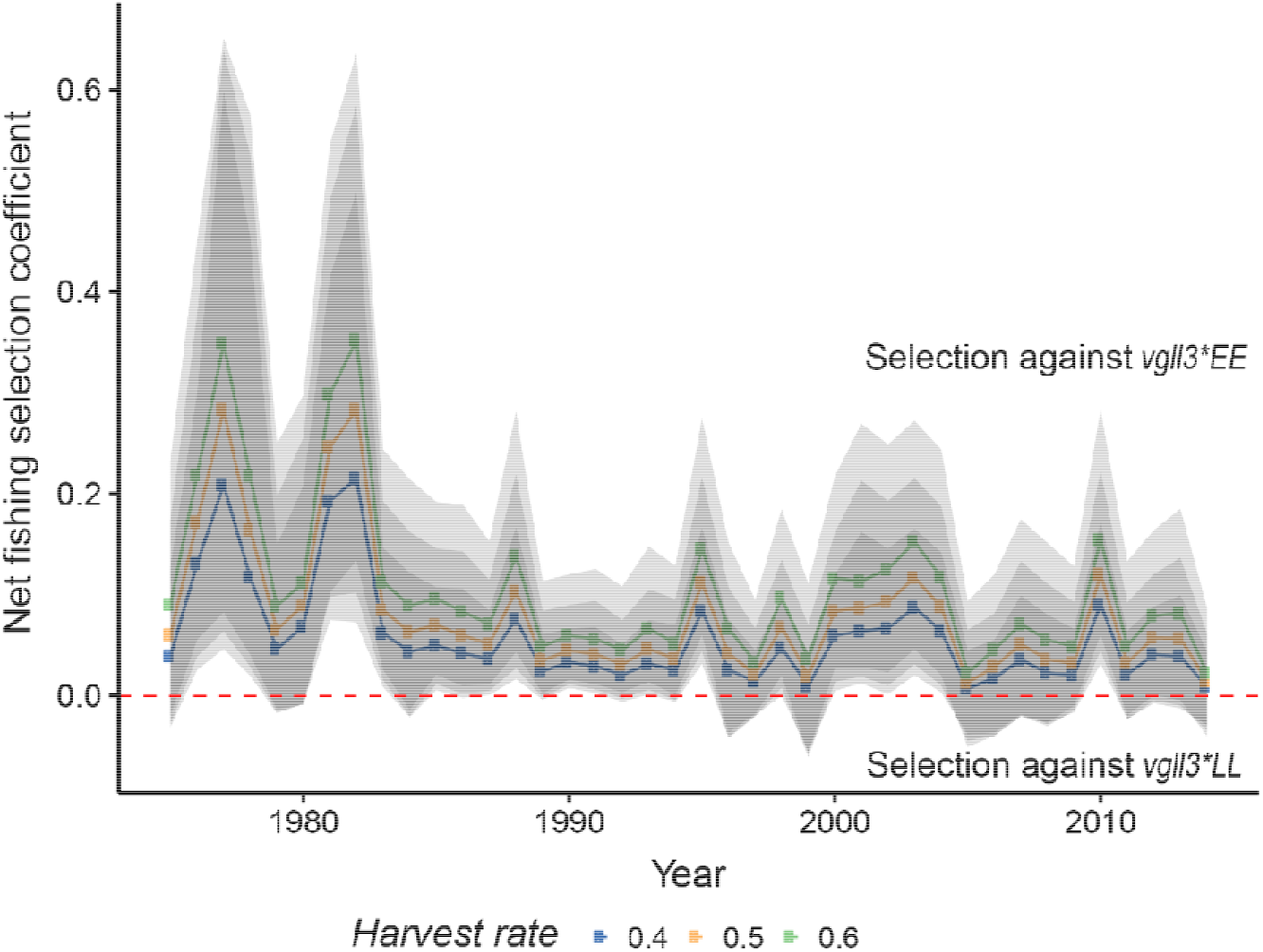
Predicted variation of net fishing induced selection over 40 years in Tenojoki as a function of harvest rate. The selection coefficient corresponds to the relative difference in survival between *vgll3*EE* and *vgll3*LL*. Calculations are based on capture probabilities (with harvest rates ranging from 40-60% of returning individuals) as a function of length, using sonar data in 2018 and 2019. It accounts for the sex-specific *vgll3* association with length during the study period, temporal variation in sex-ratio and relative use of the weir, driftnet and gillnet (see methods). Shaded area represents 95% confidence intervals based on 6000 iterations.

We provide an example of fisheries induced evolution, where both direct and indirect fishing effects contributed. Minimizing fisheries induced evolution is generally recommended as adaptation of populations to fishing can hinder adaptation to their natural environment and be costly in the long run ^21–23^. However, strong socio-economic pressures may limit opportunities for dramatic changes in fishing regulations to alter harvest rates to minimise fisheries induced evolution ^23^. In the Teno river valley for instance, salmon fishing is both a key source of income for locals through fishing tourism, as well as the foundation of the indigenous Sámi culture and identity, and restrictive fisheries regulations can be controversial ^24^. The varying selectivity of different net types may provide a means to manage the selection pressures exerted by fishing on the different ages at maturity and genotypes e.g. by regulating the relative use of the different fishing gears. Collection of additional data (e.g. additional years of genetic stock assignment and sonar counting) and the development of models coupling population dynamics and evolutionary genetics would be required prior to implementation. Such models would also help in identifying constant selective pressures that couldn’t be detected using regressions and in determining the impact of fisheries induced selection on population growth rate. Commercial harvesting of an important salmon prey species, capelin, indirectly induced evolution of Atlantic salmon age at maturity toward younger, smaller individuals. Several studies have noted the potential for a fishery to induce ecological effects beyond the target species ^25^. However, evolutionary effects have not been demonstrated earlier ^5,13^. This case also has important implications of a more applied nature. About 90% of the forage fish catch are used for fish oil and fishmeal to feed farmed animals ^6^ and salmonid fish aquaculture is the 4^th^ highest consumer ^26^. For instance in 2012, 75,800 tons of capelin was used in Norwegian aquaculture salmon feed (representing 15% of the marine ingredients ^27^). Our results therefore identify yet another indirect path by which Atlantic salmon aquaculture can negatively affect wild populations of the same species and emphasize the importance of identifying alternative, sustainable, protein sources for the aquaculture industry.

## Material &methods

### Sampling, genotyping and population assignment

The Teno river (Deatnu in Sámi, Tana in Norwegian), located at the border between Finland and Norway and draining into the Barents Sea (68 − 70°N, 25-27°E), hosts large and diverse Atlantic salmon populations ^20,28^. Scales of more than 150 000 salmon caught in different parts of the Teno river system have been collected since 1971, along with individual information including sex, length, weight, sea age at maturity, fishing gear (driftnet, weir, gillnet or rod) and fishing location. Of these, around 113 000 were collected in the mainstem, the focal region of this study. Teno salmon can remain from 2 to 8 years in fresh water before migrating to the sea. They then stay in the marine environment, where most growth occurs, for mostly 1 to 3 years, occasionally 4 or 5, before maturing and returning to spawn ^20^. Details about sample collection, DNA extraction, genotyping (at 191 SNPs, including *vgll3*, and a sex marker, sdy ^29^) and population assignment are described in ^3^. Briefly, a total of 1319 individuals genetically assigned to the Tenojoki population in the middle reaches of the mainstem of the river system (hereafter Tenojoki) in ^3^ were used in this study, resulting in an average annual sample size of 33 individuals per year between 1975 and 2014.

### Environmental variables

Evolution of age at maturity can be driven by factors occurring either in the freshwater or marine phases of salmon life history. Changes in pelagic communities of the Barents Sea may affect Atlantic salmon survival and life history traits during the marine feeding phase e.g. ^30,31^. The stock biomass of important salmon prey species (capelin, herring and krill) have been monitored for decades in the Barents Sea. Data about krill biomass (1980 – 2013) were taken from ^32,33^. Capelin biomass estimated from acoustic survey and the landed capelin catches were derived from ^34^ for 1973 – 2013. Herring dynamics has been estimated using a virtual population analysis (VPA) over the last decades ^35^. Because herring migrate out of the Barents Sea once they mature, at the age of 3-5 years ^36^, only the biomass of 1-2 year old herring were used here e.g. ^17^. These herring biomass data were retrieved from ^35^ for the 1973-1998 period. Herring biomass was calculated from the number of 1-2 year old herring and the mean weight per age as reported in ^34^ for 1988 – 2013. The Pearson correlation between biomasses from those two datasets of herring was 0.99 for the overlapping period, despite values in ^34^ being lower by a factor 1.46. The data from ^35^ were standardised accordingly by dividing them by 1.46. Data about several species interacting with salmon prey were also used in the multi-species model (see below). The annual biomass of cod (a predator of forage fish) was derived from VPA analyses (^37^, table 3.24). Landed cod biomass was also taken from ^37^. Further, an index for mesozooplankton (a forage fish food source) corresponding to the sum of *Calanus* biomass indices from different parts of the Barents Sea was used ^38^. Marine temperature has often been associated with growth, age at maturity or survival in Atlantic salmon ^39,e.g. 40^. The (monthly averaged) sea temperature in the Kola section measured in the upper 200 meters was therefore included (from Pinro.com,^41^).

Salmon fisheries targeting adult salmon returning from their marine migration occur both in the coastal areas around the outlet of the Teno River (i.e. the Finnmark region), and in the river itself using fishing gears with varying degrees of selectivity on size, and hence, age at maturity ^20,42^. The total number of nets used to catch salmon in the Finnmark coastal region was calculated for each year using data from ^43^ whereas the annual number of net fishing licenses, corrected for the number of fishing days allowed per week (which changed from four days to three days in 1980) was used as a surrogate for riverine net fishing effort. The corrected number of net fishing licenses had an among years Pearson’s correlation of 0.75 (t_34_ = 7.35, P-value < 0.001) with the number of weirs in the Teno river, estimated from count data during the 1976-2014 time period and adjusted for the number of fishing days in a week (data from years 1980 to 1982 were missing for the latter).

A sonar count provided an estimate of the size distribution of ascending salmon, thus enabling an estimation of the size selectivity of riverine fishing methods via comparison with the size distribution of salmon caught with each fishing gear type during the fishing period in June and July. An ARIS explorer 1200 sonar unit (Sound Metrics Corp., Bellevue, Washington, USA) was placed c. 55 km upstream of the Teno river mouth in 2018 and 2019. Only individuals with a length greater than 43 cm were considered as salmon, because of the occurrence of other fish species which are mostly smaller than salmon. The number of upstream migrating salmon was calculated as the difference between the number of ascending and descending individuals. In 2019, the total number of salmon in the 136 cm size group was adjusted from minus one (meaning overall, one more fish in the size group descended than ascended), to zero. The length of salmon caught in 2018-2019 with driftnet, weir, gillnet and rod was recorded by fishers (N = 17, 745, 95 and 219 in 2018 and N = 45, 717, 71 and 334 in 2019; respectively). Salmon lengths from sonar data and catches were grouped in 2 cm bins from 44 cm to 151 cm and used for estimating selectivity of different fishing methods (see below). A small number of individuals that were estimated to be larger than 150 cm in the 2018 sonar count (N = 17) were grouped with the last size class bin. One 136cm rod-caught individual was moved to the 134-135cm bin in 2019 as no individual of that size class was observed by sonar in that year.

### Statistical analyses

#### Driver of vgll3 evolution

To identify the environmental drivers of *vgll3* evolution, the proportion of *vgll3*L* alleles (0, 0.5 or 1) in an individual was regressed using a Generalized Linear Model (GLM) with the quasi-binomial family e.g. ^44^. The prey biomass (krill, capelin and herring) and sea temperature were averaged over the year(s) each individual spent in the marine environment, which was inferred from scale growth ring information. The logarithm of each prey’s biomass was included as an independent variable to allow non-linear associations with the *vgll3* proportions and a better correspondence with the multispecies model described below. The mean sea temperature, the number of coastal fishing nets, the number of tourist rod-fishing licenses (days^-1^) and net-fishing licenses in the Teno river, corrected for the number of permitted weekly fishing days, were also included as predictors. Model selection was performed with a hypothesis testing approach using backward selection with F tests. Statistical tests throughout the study were two-tailed, using an alpha value 0.05 and were performed using R^45^. The AICc of all possible models were also calculated along with the relative importance of the different variables, using the MuMIn package ^46^. Model averaging with the classical method ^47^ was performed on the subset of models accumulating an AICc weight of 0.95. For graphical representations, fitted allele frequencies were averaged per hatching year and bias-corrected and accelerated bootstrap intervals (BCa) were calculated with the BOOT package ^48^.

Co-occurrence of temporal trends in both the dependent and independent variables may lead to spurious detection of environmental effects. To account for this, de-trending of the data was achieved as advised in ^49^ by conducting residual regression on the original data ^15^. To do this, the independent variables included in the GLM were regressed in a linear model against years and squared years to estimate linear or quadratic trends, respectively. The squared year term was then removed from the model if not significant (assessed by F tests). Residuals of these models then replaced the original independent variables. The GLM described above was re-run by including an additional spawning year effect to remove temporal trends in *vgll3* allele frequency. The spawning year was kept as a covariate during the model selection process even if it was not significant.

#### Drivers of sex-ratio changes

To identify variables modifying the sex-ratio of returning adults in Tenojoki, a binomial GLM was performed by coding the sex of individuals as 0 (male) or 1 (female). The log biomass of prey species, sea temperature, number of coastal fishing nets and number of riverine net and rod (days^-1^) fishing licences were included as independent variables as previously. Model selection was realized using the Likelihood ratio test (backward variable selection) and the AICc criterion. This analysis was repeated by using the residual regression method to account for potential co-occurrences of temporal trends in the dependent and independent variables, as described above.

#### Drivers of changes in age at maturity

Age at maturity of individual salmon was regressed in a multinomial model using all the independent variables described above (R package nnet ^50^). Additionally, *vgll3* genotypes and sex were included in the model in a two-way interaction, due to known *vgll3* sex-specific effects ^4^. Two individuals that matured after five years at sea were considered as individuals having matured after four years at sea. Variable selection with F tests and AICc model averaging were also performed. This analysis was repeated by using the residual regression method.

#### Indirect evolutionary effects

The indirect effect of capelin harvesting on *vgll3* evolution was estimated using the Monte Carlo Method for Assessing Mediation (MCMAM)^51^ with capelin (log) biomass as the mediator. The effect of (log) capelin biomass on *vgll3*L* odd-ratio was obtained with the quasi-binomial GLM model described above (after backward selection, including the capelin, herring, krill and riverine net fishing licenses variables). The estimated parameter and standard error were used to generate 3000 samples drawn from a normal distribution. The posterior distribution of the direct effect of harvest rate on the (log) capelin biomass was obtained with the multispecies Gompertz model described below (3 000 MCMC samples, Extended Data Fig. 4, equation 1). The indirect effect of capelin harvest on *vgll3*L* odd-ratio corresponds to the product of the two sets of samples (i.e. effect of capelin fishing on (log) capelin x effect of (log) capelin on *vgll3* odds) and thus accounts for uncertainty in both estimates. The indirect effect was also evaluated graphically to account for more than one interaction, by:

1. Using the multispecies Gompertz model to predict the biomass dynamics of capelin, herring and krill when capelin harvesting is set to zero from the first year of the time series;
2. Using the newly generated data (posterior median) to predict the *vgll3* allele frequency variation using the quasi-binomial GLM obtained with backward selection
3. Plotting the temporal allele frequency variation with Bca confidence intervals

This analysis accounted for the effect of capelin harvesting on the biomass of all the Gompertz model species but didn’t account for *vgll3* evolutionary dynamics (i.e. the propagation of the indirect negative effect from one generation to the next).

#### Multispecies Gompertz model

Fish populations in the Barents Sea have experienced large variation in their abundance over the last 40 years due to fishing and predation e.g. ^16^. Consequently, important indirect effects on Atlantic salmon age at maturity variation/evolution may be expected. In order to estimate potential indirect effects, the Barents Sea species interactions, the effect of temperature, density dependence and of cod and capelin fishery were estimated using the multispecies Bayesian Gompertz model developed in ^17^. To summarize, on the log scale, the process equations were:

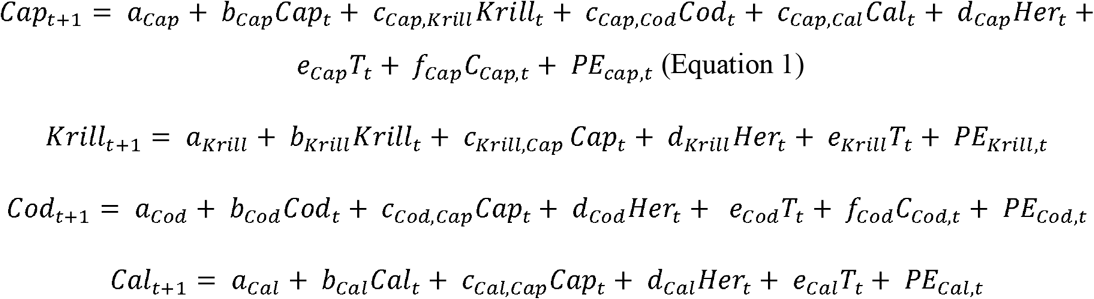

With Cap_t_, Krill_t_, Cod_t_, Cal_t_ and Her_t_ being the log biomass of capelin, krill, cod, *Calanus* and herring, respectively, in year *t*, T_t_ the annual sea temperature in the Kola section of the Barents Sea at 0-200 m depth (along the 33°30’E meridian from 70°30’ to 72°30’N) in year *t* and C_Cap,t_ and C_Cod,t_ the capelin and cod harvest rates, respectively, in year *t* (catch/annual biomass). The coefficients *a, b* and *c* represent the productivity, density dependence and interactions among modelled species, respectively. The coefficient *d* represents the interactions with herring, *e* the temperature effects, *f* the fishing effects and *PEs* the multivariate-normal-distributed process errors. The prior distributions in the model were identical to ^17^ (e.g. uniform prior distributions from −10 to 10 for all process parameters *a, b, c, d, e* and *f*).

Posterior distributions were approximated using MCMC methods with the Just Another Gibbs Sampler software JAGS ^52^. The model was run for 1 410 000 iterations including a burn-in length of 710 000. One iteration out of 1000 was kept and 3 chains were run. Convergence was assessed using the Gelman and Rubin’s convergence diagnostic and a Potential Scale Reduction Factor (PSRF) threshold of 1.15 ^53^. The model posterior medians were substituted to the original data for other analyses, in order to replace missing values in krill from 1973 to 1979 and employ common data between the different models used for indirect effect evaluations.

#### Net fishing size selectivity

Teno local net fishing encompass three fishing methods with different size selectivities ^20^. To better understand the potential effect of net fishing on *vgll3* allele frequency changes, fishing size selectivity in Tenojoki salmon was estimated by using sonar count data and the length distribution of catches per fishing gear in 2018 and 2019 (including driftnet, weir, gillnet and rod). The number of salmon caught per 2 cm. length class (*l*, from 44 to 151 cm) was analyzed for each gear (g) with a Generalized Additive Model (GAM) using the beta-binomial family to account for overdispersion. Length of each salmon (*L*) was included as a predictor, in a regression spline (*s*_*g*_):

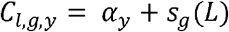

With *c*_*l,g,y*_ the mean proportion of salmon caught per 2 cm. length class (*l*) in year (*y*) with the fishing gear (g) and, *α*_*y*_ the annual intercepts with priors following a normal distribution (mean = 0, sd = 10). The dispersion parameters of the beta distribution (i.e. addition of both shape parameters) followed a prior uniform distribution between 2.5 and 5000 for each gear. The priors, code and associated data for the smooth part of the model were generated with the R package *mgcv* (see ^54^ for more information). The model was run in JAGS ^52^ (R package runjags ^55^) for 7 600 000 iterations including a burn-in period of 4 000 000. Two chains were run and one iteration out of 1200 was kept. Convergence was checked using the Gelman and Rubin’s convergence diagnostic ^53^ and inspection of trace plots. *C2*_*i,l,g*_, used below, corresponds to the mean capture probability of the gear (*g*) per 2 cm. length class (*l*) calculated at each iteration (*i*) from the model estimates, and divided by 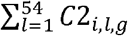 so that it sums to one over the different length groups.

#### Temporal variation in net fishing selection on vgll3

The effect of riverine net fishing on *vgll3* allele frequency depends on the size selectivity of the fishing gears, calculated above, the harvest rate per fishing gear and distribution of *vgll3* genotypes according to salmon size. The number of salmon assigned to the mainstem population was modelled with a GAM using the negative binomial family, to allow for over-dispersion. Length was included as an independent variable inside annual random smooths, to account for changes in age at maturity over time. All GAM models in the following part of the method were run using the mgcv package^56^. To account for uncertainty, simulations from the posterior distribution of the fitted model were performed by using a Metropolis Hastings sampler ^57^ (8 chains, 300 000 iterations per chain, thinning of 400, burn-in of 1000). For each iteration (*i*), the proportion of salmon in the different 2 cm length classes (*Pl*_*i,l,y*_) was calculated for each year (*y*) from the model predicted values. Those proportions were later used to estimate the gear specific fishing mortality to match the expected gear-specific annual harvest rates (see below). A model including annual random smooths per sex was also used to estimate *Pl2*_*i,l,s,y*_, the proportion of salmon in the different 2cm length class per sex and year at each iteration (*i*) (Metropolis Hastings sampling with 8 chains, 300 000 iterations per chain, thinning of 400, burn-in of 1000).

The proportion of salmon caught by each fishing gear over the 40-year period (*Pr*_*y,g*_) was estimated from the annual catch weight, converted to an estimate of the individual number using the annual average weights of salmon caught in the mainstem with the different gears (N = 84 452). The total harvest rate (*HR*_*tot*_) was fixed at either 40%, 50% or 60% (as riverine fishing mortality of up to 69% on tagged salmon has been reported^58^), only the gear specific harvest rate 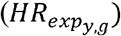 being allowed to vary over time according to the previously calculated proportions (*Pr*_*y,g*_):

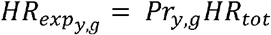

Given that fishing with the different gears occurs simultaneously, the instantaneous fishing mortality needed to be estimated according to the expected harvest rate 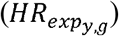 and gear size selectivity (*C2*_*i,l,g*_). A modified version of the Baranov’s catch equation was used to calculate the harvest rates (*HR*_*i,y,g*_) of the gear (*g*) in year (*y*) at each iteration (*i*):

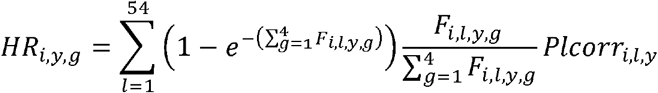

With *Plcorr*_*i,l,y*_ the proportion of salmon in the 2 cm. length class (*l*) in year (*y*) at iteration (*i*) before fishing (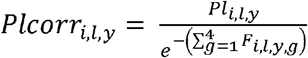, modified to sum to one over the length classes). *F* _*i,l,y,g*_ is the instantaneous fishing mortality for the length class (*l*), gear (*g*) and year (*y*) calculated at each iteration from the mean capture probabilities *C2*_*i,l,g*_:

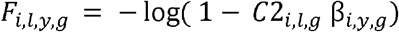

β_*i,y,g*_ was determined by minimizing 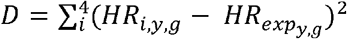 using the R optim function from the stats r package with the “L-BFGS-B” method ^59^ and a lower bound of 0.01. The Nelder-Mead method was used if convergence was not achieved with the “L-BFGS-B” method (12 iterations), while ensuring that β_*i,y,g*_ was positive (e.g. by squaring it). The maximum instantaneous fishing mortality was capped at 2.30, allowing a maximum harvest rate per fishing gear of 0.95. This was used to avoid unrealistic correction, at some iterations, of the length-frequency distribution when calculating *P1corr*_*i,l,y*_.

The exploitation rates per year (*y*), *vgll3* genotype (*v*) and net fishing gear (*g*) were then calculated at each iteration (*i*) as follows:

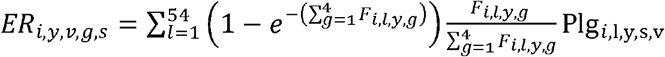

With Plg_*i*,l,y,s,v_ the proportion of individuals of sex (*s*) in the 2 cm. length class (*l*) in year (*y*) and with the *vgll3* genotype (*v*), calculated at each iteration (*i*) as follow:

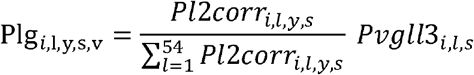

And modified to sum to one over the length class. *P12corr*_*i,l,s,y*_ is the proportion of individuals of sex (*s*) in the length class (*l*) in year (*y*) before fishing, calculated at each iteration (*i*) from the length distribution previously estimated with the GAM:

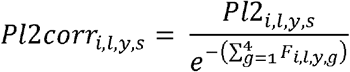

*Pvgll3*_*i,l,s*_ corresponds to the probability of the different *vgll3* genotypes for a salmon in the 2 cm length class (*l*) and sex (*s*). This probability was estimated with a multinomial GAM including the 2 cm length classes in a regression spline for each sex and genotype. Posterior simulations (1 chain, 6 000 000 iterations, thinning of 1000, burn-in of 1000) were performed using the Metropolis Hastings sampler. Alternative models allowing the smooth parameter to change per sex and decades in one or both of the modelled genotypes were also run. There was no evidence for changes in the length-specific probability of the different *vgll3* genotypes over time (model fitted with maximum likelihood, ΔAIC > 8 and LRT test P-value = 0.096 with the best competing model).

Finally, the mortality per genotype (*v*) and year (*y*) for the 3 net fishing methods in the population corresponded to:

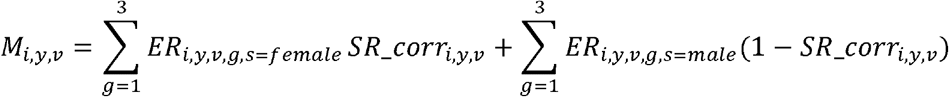

The mortality per genotype accounts for temporal variation in the proportion of females per genotype before fishing (*SR_corr*_*i,y,v*_). Temporal variation in the female salmon proportion was analysed with a quasi-binomial GAM, including year as an independent variable inside a cubic regression spline for each genotype and adding an extra penalty to each term. To account for uncertainty, simulations from the posterior distribution of the fitted models were performed by randomly drawing 6000 values from a multivariate normal distribution with the mean vector and the covariance matrix equal to the model estimates. The GAM fitted values were calculated at each iteration to obtain *SR*_*i,y,v*_. The proportion of females before fishing corresponded to:

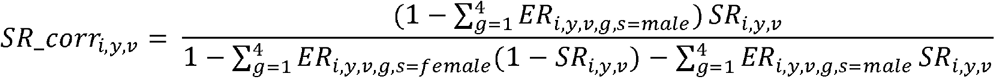

The survival of the different genotypes could then be calculated:*W* _*i,y,v*_ *=1* − *M*_*i,y,v*_. The annual fishing selection against the less fitted homozygote (*S*_*y*_) was calculated as follows:

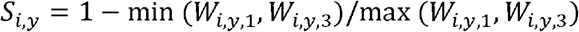

The sign of *s*_*y*_ was changed according to *max* (*W* _*i,y,1*_, *W*_*i,y,3*_); *S*_*y*_ is positive when *W*_i,y,1,_ < *W*_*i,y*,3_ and negative otherwise. Calculations of variation in selection and relative mortality assume temporally constant gear selectivity. The total harvest rate is also assumed constant over time and was included to assess whether the fishing pressure in the Teno river is strong enough to induce significant selection.

Effect of changes in net fishing selection on *vgll3*, at a constant harvest rate of 0.5, were investigated in a post-hoc analysis by included s_i,y_ in the normal and de-trended quasi-binomial GLM previously described.

## Data and code availability

The datasets used in the current study will be uploaded to a public data repository upon acceptance.

## Code availability

The personal custom codes used in the current study will be uploaded to a public data repository upon acceptance.

## Availability of biological material

All unique biological materials used are available from the authors

## Acknowledgements

We thank fishers who collected scales and phenotypic information and people who organized and read the scales, especially Jorma Kuusela and Jari Haantie, as well as Jan-Peter Pohjola for sonar data reading, and Maija Länsman for compiling the detailed river catch and effort data. We also thank Eva Eriksen for providing corrected krill biomass data and Øystein Langangen for sharing codes and information on the multispecies Gompertz model. This project received funding from the Academy of Finland (projects No. 284941, 286334, 307593, 302873, 318939 and, 325964) as well as from the European Research Council (ERC) under the European Union’s Horizon 2020 research and innovation programme (grant agreement No 742312). Part of YC salary was funded by the Norwegian Research Council (projects No. 275862 EcoEvoGene and 280308 SeaSalar).

## Author contributions

J.E, and P.O. coordinated the collection of scale samples and river fisheries data; C.R.P., Y.C., T.A. and J.E. designed the study; Y.C realized the lab work, Y.C. analysed the data; Y.C., C.R.P. and

T.A. wrote the manuscript and all authors contributed to its revision.

## Materials &Correspondence

Correspondence and requests for materials should be addressed to C.R.P.

## Extended Data

**Extended Data Figure 1:**
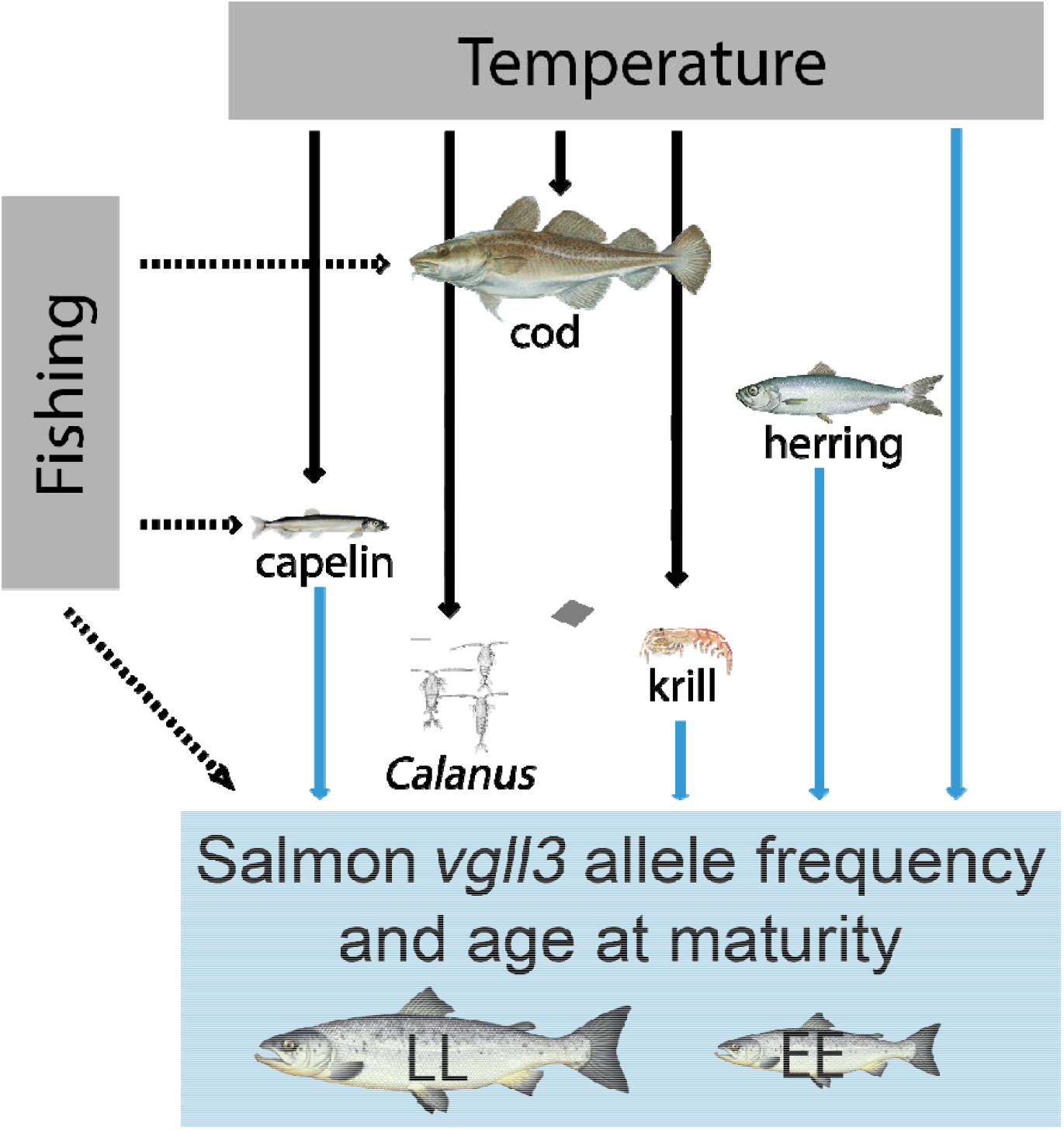
Interactions in the Barents sea community as modelled by ^17^ and links with Atlantic salmon age at maturity and *vgll3* allele frequencies.

**Extended Data Figure 2:**
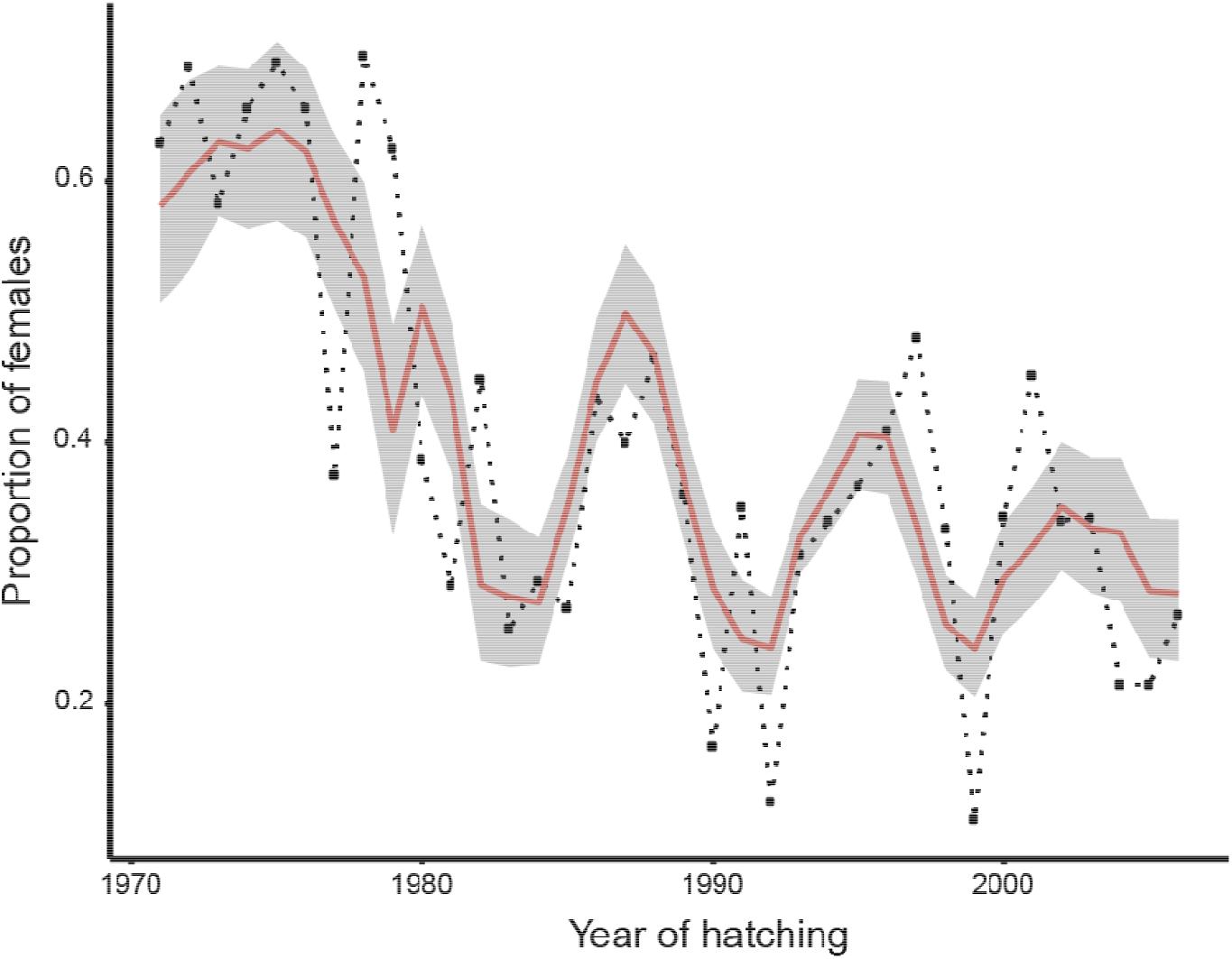
Temporal changes in proportion of females over 36 hatching years. The black dotted line represents the observed data, the red line the female proportion averaged from individual fitted values. Confidence intervals are bias-corrected and accelerated (BCa) 95% bootstrap interval based on 3000 replicates.

**Extended Data Figure 3:**
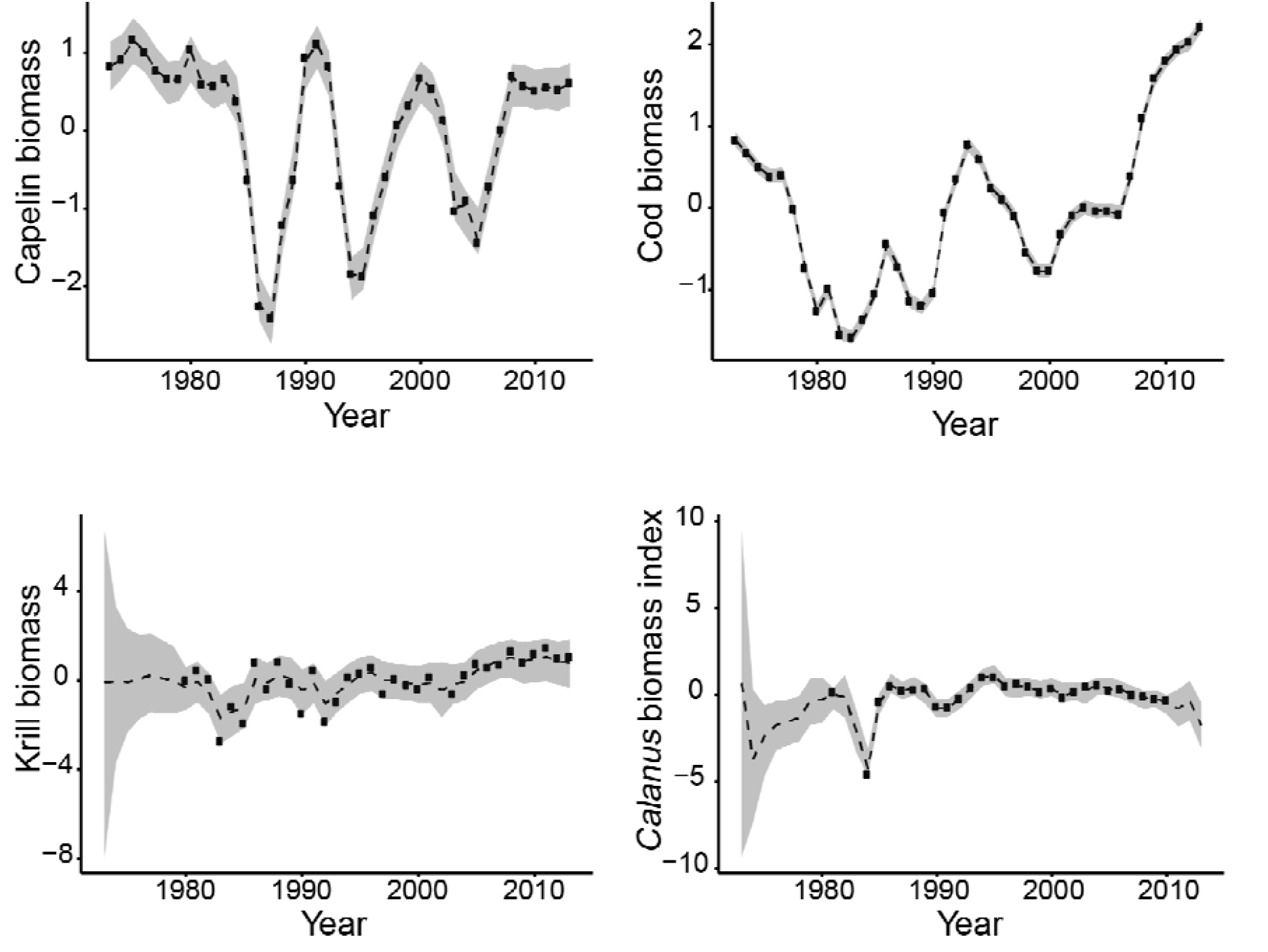
Annual variation in log transformed and normalized biomass of capelin, cod, krill and, *Calanus* in the Barents Sea. The dots represent the observed data, the dashed lines the model posterior medians and the shaded areas the 95% credible intervals.

**Extended Data Figure 4:**
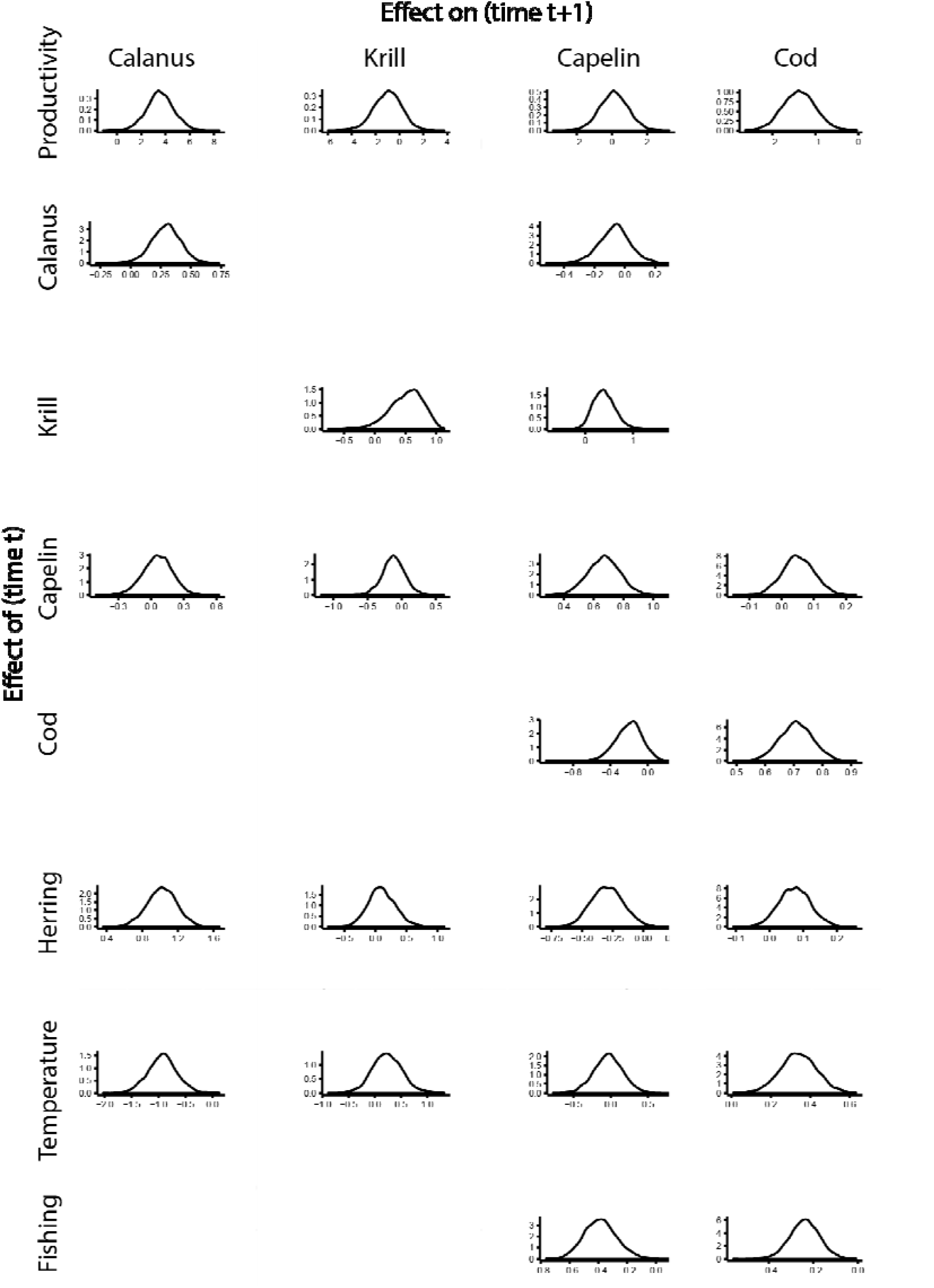
Posterior probability distribution of process parameters included in the multispecies Gompertz model.

**Extended Data Figure 5:**
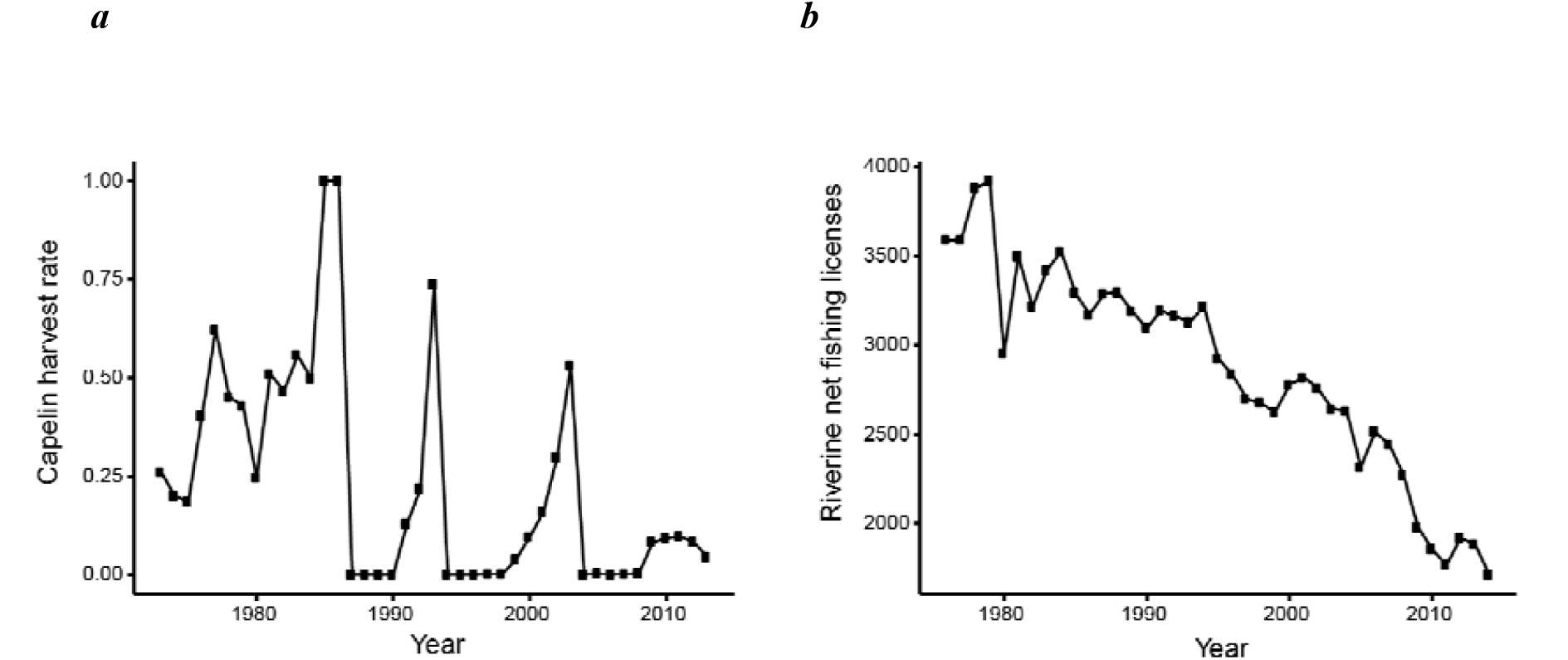
Temporal variation in fishing related indices. *a*, capelin harvest rate. *b*, corrected riverine net fishing licenses (number of licences multiplied by effective number of fishing day per week).

**Extended Data Figure 6:**
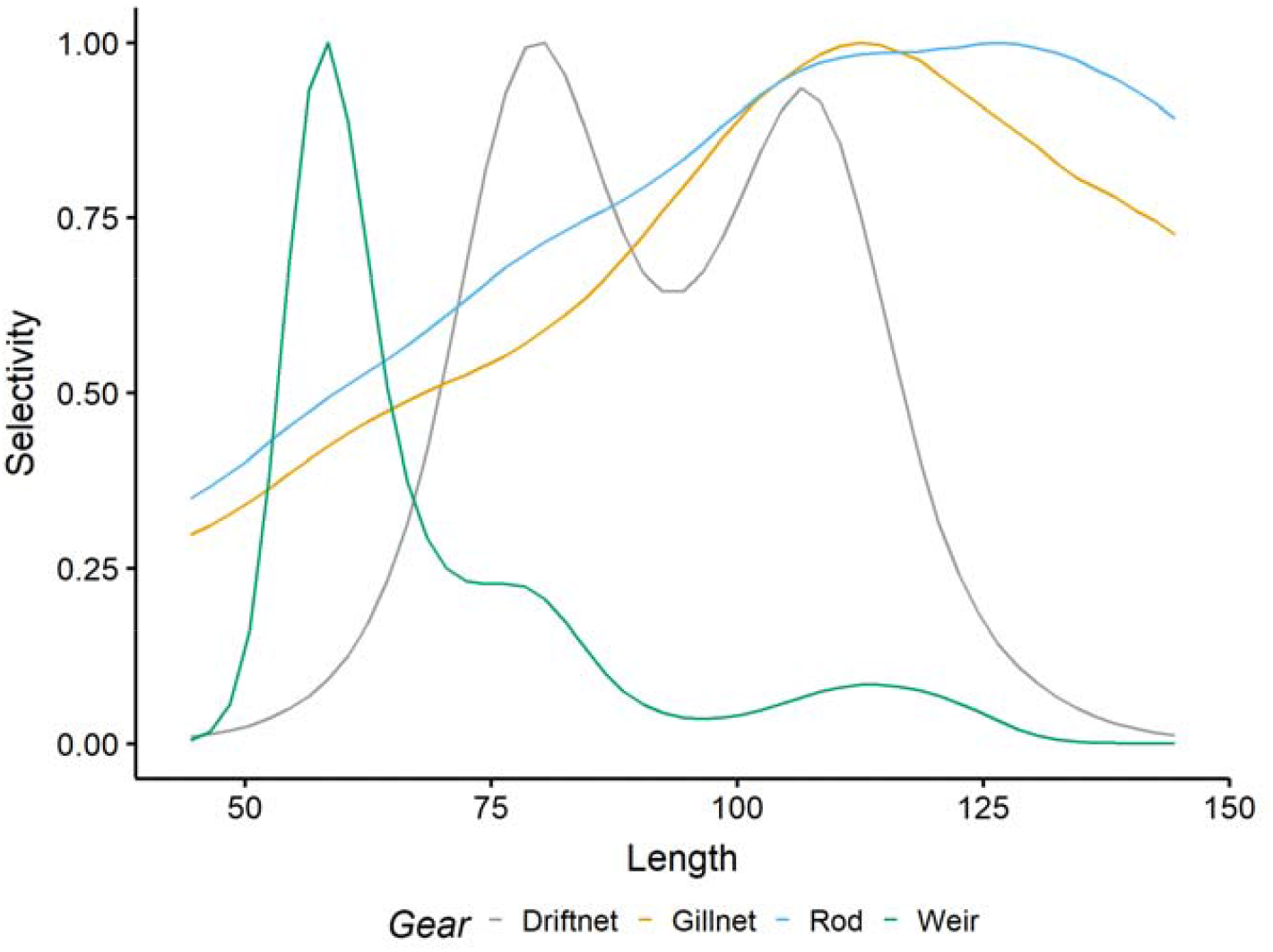
Bayesian posterior median selectivity of the different gears as a function of salmon length (in cm). Median capture probabilities were transformed so that selectivity curves peak at 1.

**Extended Data Figure 8:**
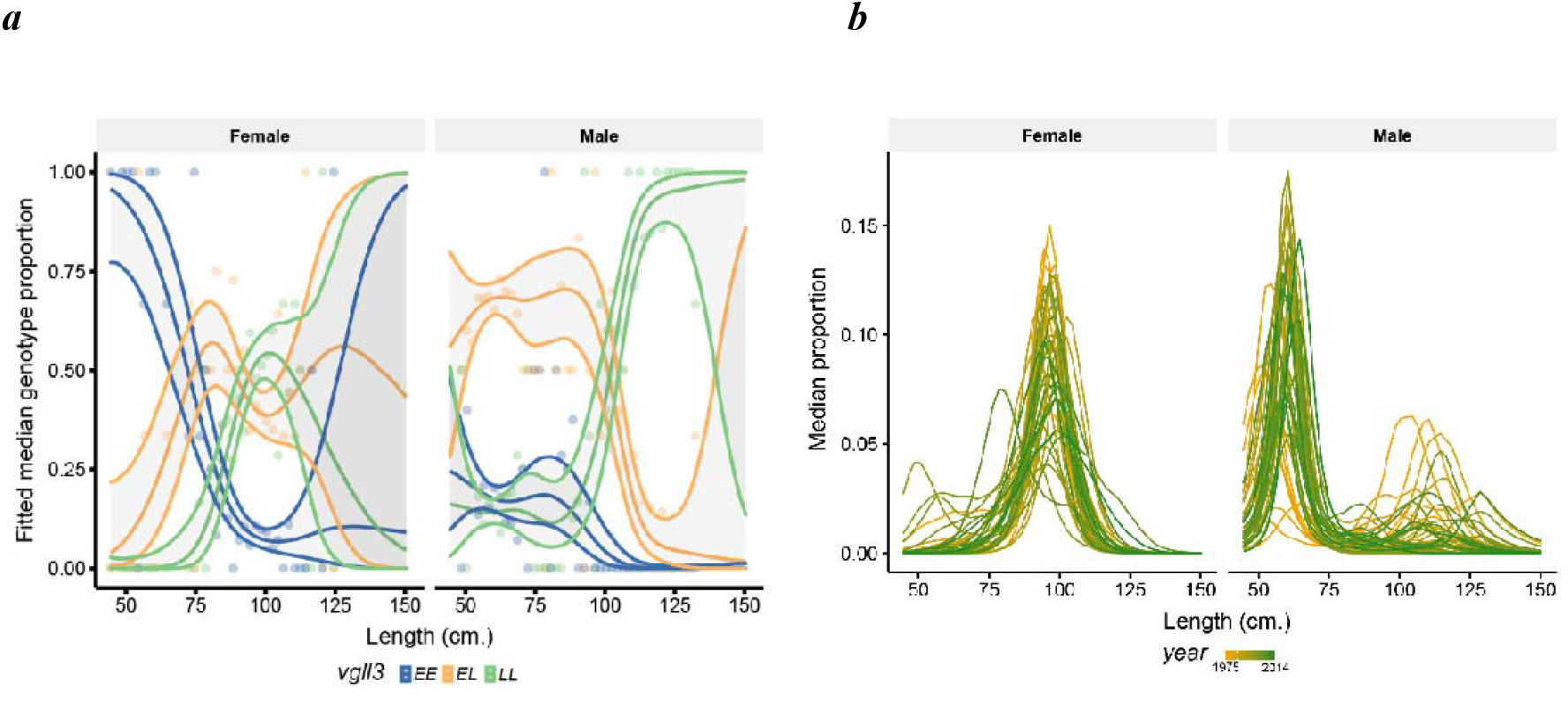
Median proportion of *a, vgll3* genotypes per length class and sex and *b*, salmon per length class, sex and year. Median proportions were calculated using metropolis Hastings sampling from **a**, a multinomial GAM and **b**, a negative binomial GAM (6000 iterations kept). Shaded area represents 95% confidence intervals.

**Extended Data Figure 7:**
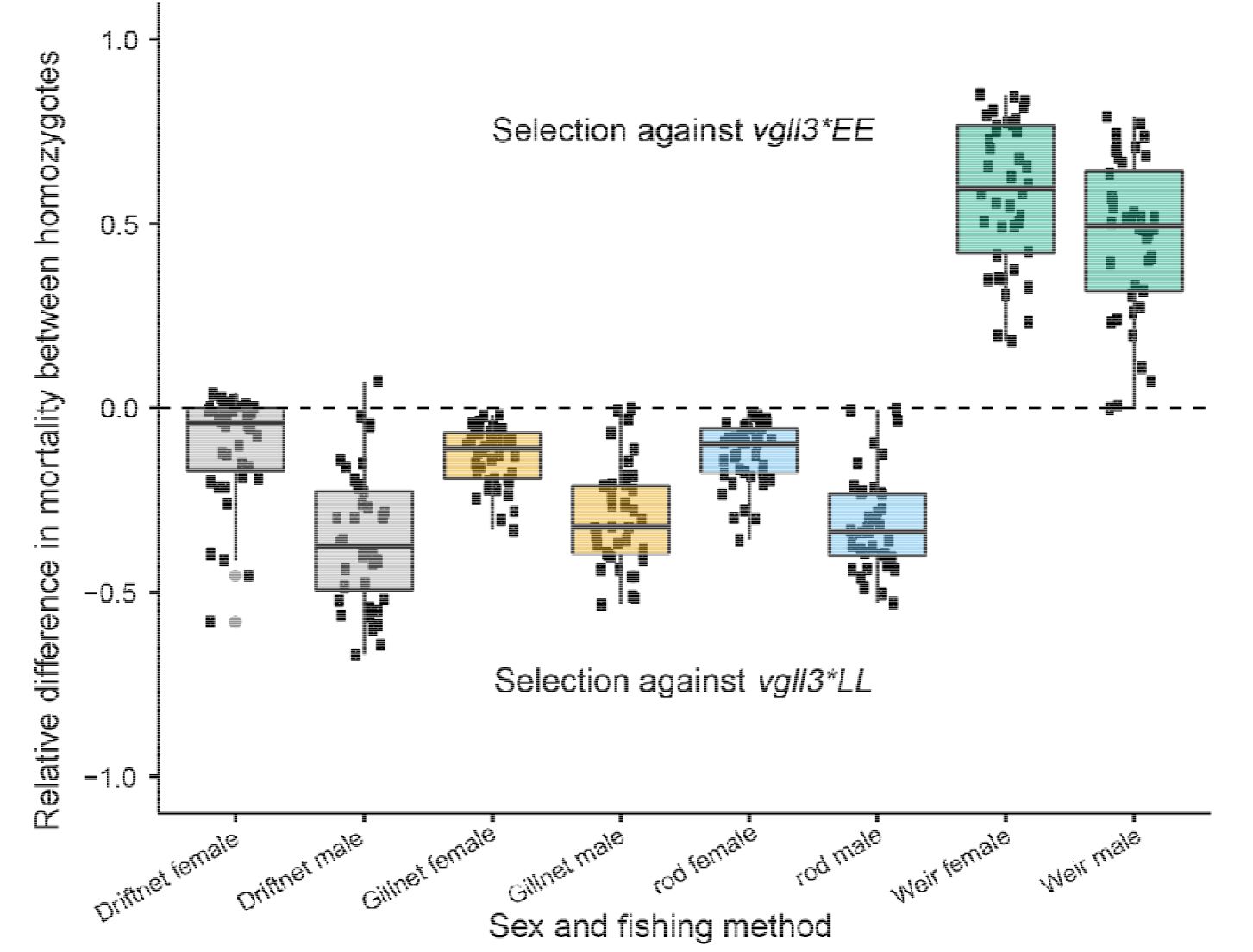
Median annual relative difference in mortality for different fishing gears in males and females. Positive values indicate higher mortality in the *vgll3*EE* genotype than in the *vgll3*LL* genotype, negative values represent higher mortalities in the *vgll3*LL* genotype. Boxplots display the minimum value (median – 1.5 interquartile range), 25^th^ percentile, median, 75^th^ percentile and the maximum value (median + 1.5 interquartile range).

**Extended Data Table 1:**
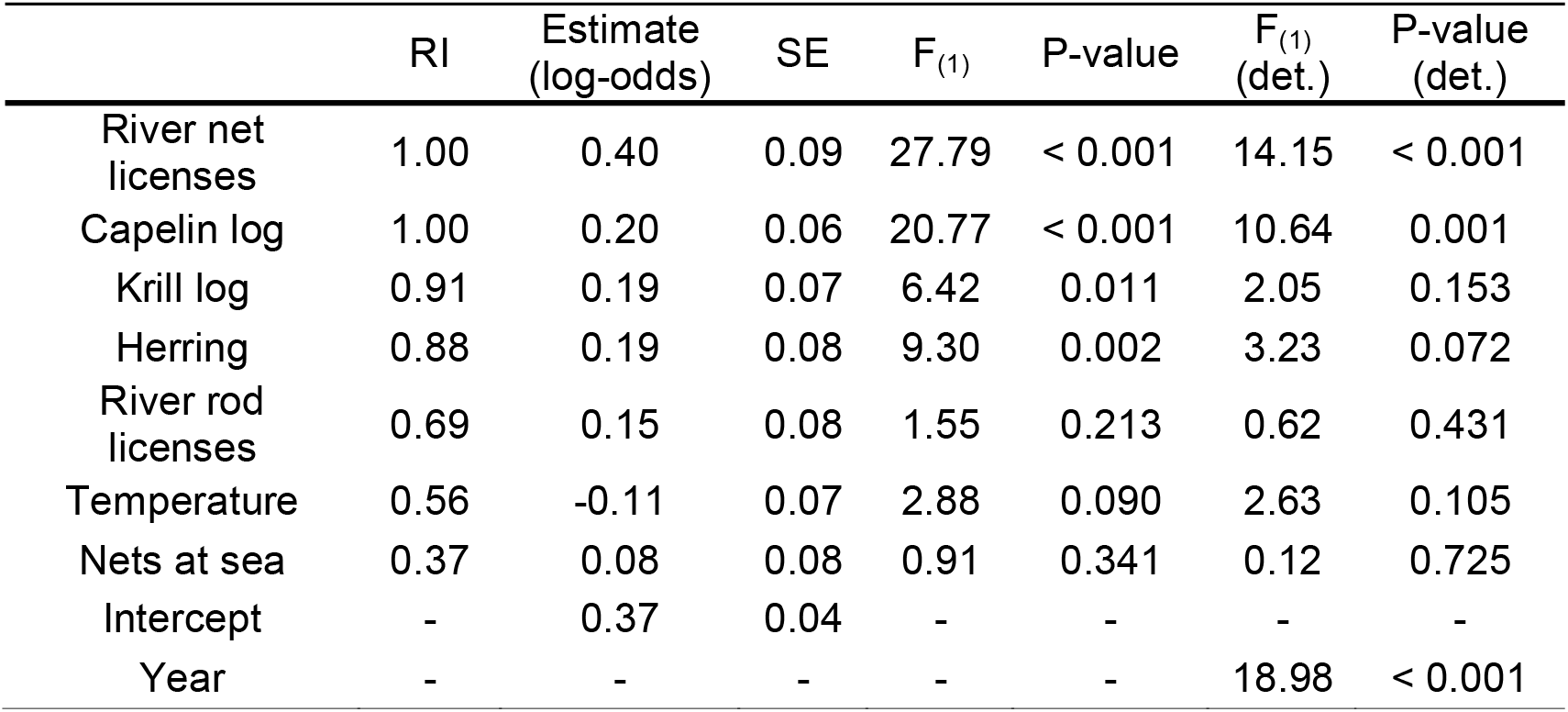
AICc relative importance (RI), model averaged standardized estimates, unconditional standard errors (SE), and hypothesis testing F-tests for the 7 predictors included in the *vgll3* allele frequency (quasi)binomial regressions. Det. - de-trended.

**Extended Data Table 2:**
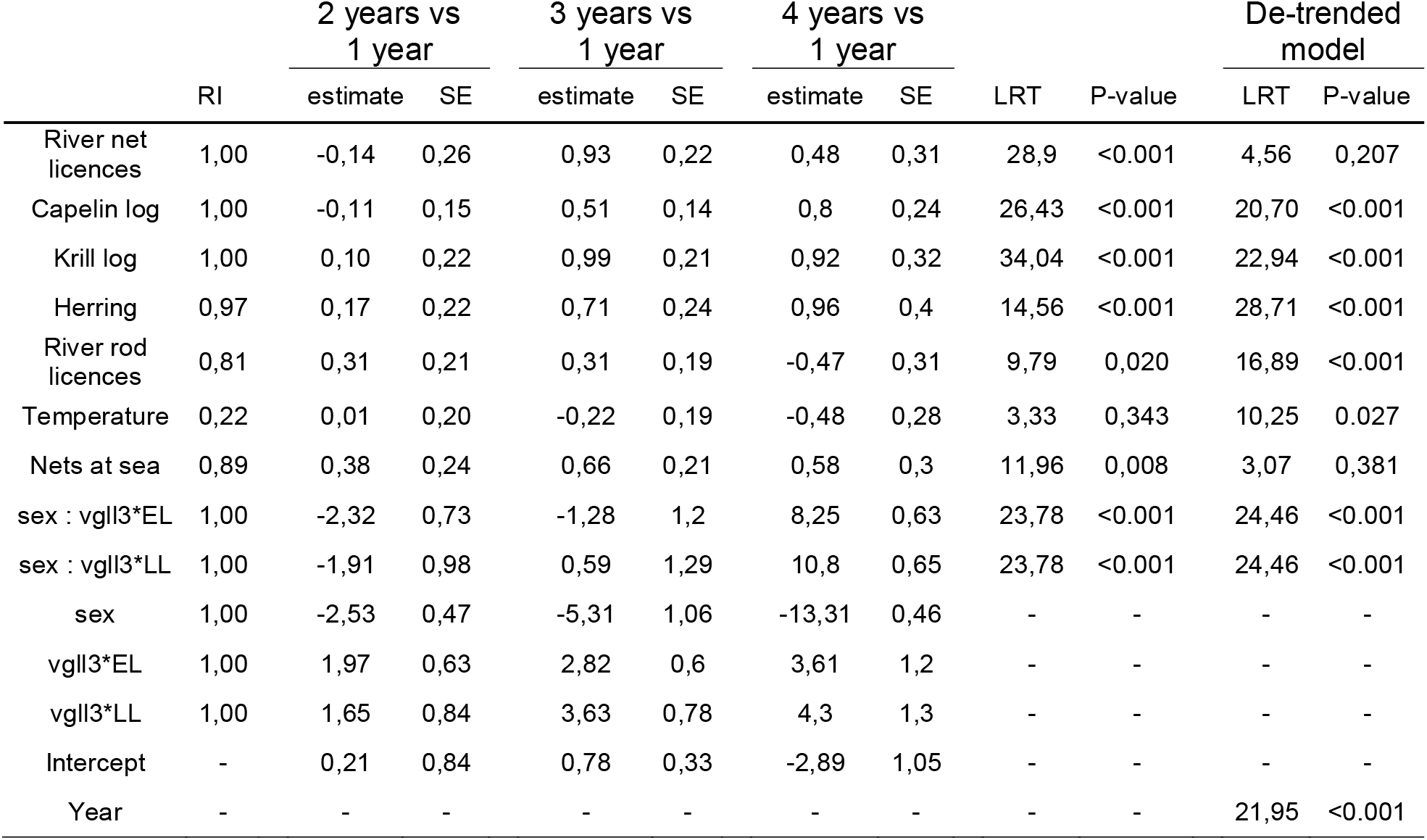
AICc relative importance (RI), model averaged standardized estimates, unconditional standard errors (SE), and hypothesis testing LRT-tests for the 9 predictors potentially influencing age at maturity probability in salmon individuals in the dataset (multinomial model).

**Extended Data Table 3:** AICc relative importance (RI), model averaged standardized estimates, unconditional standard errors (SE) and hypothesis testing LRT-tests for the 7 predictors included in the binomial regressions for the probability to sample a female. Det. for de-trended.

|  | RI | Estimate<br>(log-odds) | SE | LRT<sub>(1)</sub> | P-value | LRT<sub>(1)</sub><br>(det.) | P-value<br>(det.) |
| --- | --- | --- | --- | --- | --- | --- | --- |
| River net licences | 1.00 | 0.52 | 0.12 | 22.32 | < 0.001 | 0.35 | 0.553 |
| Capelin log | 1.00 | 0.40 | 0.07 | 34.06 | < 0.001 | 14.28 | < 0.001 |
| Krill log | 1.00 | 0.44 | 0.11 | 15.94 | < 0.001 | 10.79 | 0.001 |
| Herring | 1.00 | 0.50 | 0.11 | 20.84 | < 0.001 | 24.86 | < 0.001 |
| River rod licences | 0.30 | 0.07 | 0.11 | 0.55 | 0.458 | 4.53 | 0.033 |
| Temperature | 0.32 | -0.07 | 0.10 | 0.42 | 0.517 | 1.70 | 0.192 |
| Nets at sea | 1.00 | 0.51 | 0.11 | 25.20 | < 0.001 | 16.78 | < 0.001 |
| Intercept | - | -0.48 | 0.06 | - | - | - | - |
| Year | - | - | - | - | - | 37.27 | < 0.001 |

## Supplementary notes

### Drivers of *vgll3* allele frequency changes

The maximum variance inflation factor calculated, for the number of nets at sea, was 3.79, below the frequently recommended thresholds. The top ranked model based on the AICc had an Akaike weight of 0.22, indicating model uncertainty. It included the number of riverine net and rod licenses, capelin, krill, herring (log) biomasses and temperature covariates. The number of riverine net licenses, capelin, krill and herring were all included in the 5 other models within a ΔAICc inferior to two. Model parameters were averaged with the conditional method over 16 models having a summed weight of 0.95 (Extended data Table 1). The dispersion parameter was 0.89 in the full model.

### Genetic and plastic basis of phenotypic changes

Using a multinomial regression, we investigated the role of the environmental variables on the probability to observe the different age at maturity classes. After controlling for the sex-specific genetic effects, we found that the variables positively associated with *vgll3*L* were also positively associated with the probability to observe older, later maturing Atlantic salmon (Extended Data Table 2). The total number of nets at sea was also positively associated with later maturation probabilities (χ^2^_(3)_ = 11.96, P-value = 0.008, Extended Data Table 3). However, the number of nets at sea and the number of riverine net fishing licenses were no longer significant in the de-trended regression (χ^2^_(3)_ = 3.07, P-value = 0.381, χ^2^_(1)_ = 4.56, P-value = 0.207, respectively; Extended Data Table 2). The model obtained with backward variable selection also obtained the highest AICc, with a ΔAICc superior to 3, indicating strong support. Model averaging was performed over 7 models (Extended Data Table 2).

### Proportion of females

Using a binomial model, we found that variables positively associated with the *vgll3*L* allele frequency were also positively associated to the probability to sample a female (Supplementary notes; Extended Data Table 3, Extended Data Figure 2). The number of nets at sea was also positively correlated with female proportion (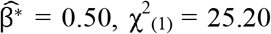, P-value < 0.001, Extended Data Table 3). Temporal variation in number of nets at sea is particularly influenced by the cessation of marine driftnet fishing in 1989, the fishing method the most used in the previous period and selecting preferentially 1SW and small 2SW ^42^, corresponding mainly to males in Tenojoki. In the de-trended model, the number of riverine net licenses was not significantly associated with female proportion anymore (Extended Data Table 3), however the number of rod fishing licenses became marginally positively associated with the probability to sample a female (Extended Data Table 3). Four models cumulating an AICc weight greater than 0.95 were used in the model averaging.

### Multispecies Gompertz model

The multispecies Gompertz model reproduces the observed variation in biomass indices well (Extended Data Fig. 3). The correlation coefficients between the observed and modelled biomass indices were greater than 0.91. The negative effect of fishing on capelin biomass (Extended Data Fig. 4) was used to quantify the indirect effect on *vgll3* allele frequency.

### Net fishing selection

#### Gear selectivity

Selectivity estimated from sonar data indicated specific patterns for each gear type (Extended Data Fig. 6). Weir selectivity was bell-shaped with a modal value of 58 cm. Driftnet selectivity was bimodal with modal values of 80 and 107 cm. Gillnet and rod had sigmoid selectivity curves with capture probabilities increasing with salmon length until reaching maxima at 112 and 126 cm, respectively.

#### Probabilities of vgll3 genotypes per length class

The probability of *vgll3* genotypes in harvested salmon differed according to the 2 cm length classes and sexes (Extended Data Fig. 8a). Small females (e.g. below 75 cm) were more likely to be *vgll3*EE* than *vgll3*EL* and *vgll3*LL*, while small males were more likely to be *vgll3*EL* and had a similar probability to be *vgll3*LL* and *vgll3*EE*.

#### Length-frequency distributions

The median length-frequency distributions of harvested salmon calculated from negative binomial GAM fitted values differed between sexes (Extended Data Fig. 8b). Females measuring around 97 cm were the most frequent whereas most frequent males measured around 60 cm. There was among-year variation in the length-frequency distribution, particularly in males, where large individuals were more frequent in the early years of the time series. The negative binomial parameter controlling the over-dispersion was 7.31.

#### Relative fishing mortality per sex and gear

The expected relative difference in fishing mortality between alternative *vgll3* homozygotes varied greatly among gears, sex and years (Extended Data Fig. 7). On average, the annual relative difference in mortality was higher in males than females for fishing gears selecting against *vgll3*LL* (i.e. driftnet, gillnet and rod) but lower for weir fishing selecting against *vgll3*EE*. Fishing mortality of *vgll3*LL* was on average 52% lower than *vgll3*EE* when weir was used but 27-29% higher when driftnet, gillnet or rod were used. The harvest rate had a negligible effect on the calculated difference in mortality between homozygotes. The annual medians calculated with a harvest rate of 0.5 differed, on average, by 1% from those calculated with harvest rates of 0.4 or 0.6.

#### Proportion of females per year and their vgll3 genotypes

The GAM smooth term estimating temporal variation in female proportion was not significant for the *EE* genotype (edf = 0.45, F = 0.05, P-value = 0.171) where the female proportion declined linearly over time. The proportion of *LL* females decreased over time in a slightly non-linear manner (edf = 1.32, F = 0.33, P-value = 0.001) whereas the proportion of *EL* females mainly declined during the 1980s and the last 7 years of the time series (edf = 6.25, F = 2.07, P-value < 0.001). The highest change in female proportion was observed for *EL* heterozygotes which was also the genotype with the largest difference in age at maturity between sexes, due to sex-specific dominance patterns. The dispersion parameter of the quasi-binomial GAM was 1.52.

#### Temporal variation in relative mortality and selection

The total harvest rate in the Teno river, fixed at plausible values (0.4, 0.5 or 0.6), would be strong enough to induce significant net fishing induced selection (i.e. differences in survival between *vgll3*EE* and *vgll3*LL* individuals; Fig. 3). The annual variation in net selection was due to changes in the sex-specific length-frequency distributions, the relative use of the different fishing gears and the sex ratio. Regardless of the harvest rate, the median selection values were all positive and indicated net fishing selection against *vgll3*EE* individuals. Confidence intervals didn’t overlap zero in 20 of the 40 years, independently of harvest rates. The median selection coefficients were on average 2.37-2.70 times higher in the first 10 years of the time series than in the last 30 years.

The median estimates of selection coefficients calculated under a harvest rate of 0.5 were included into the *vgll3* quasi-binomial model as an independent variable. The calculated fishing selection was positively associated with the *vgll3*L* allele frequency 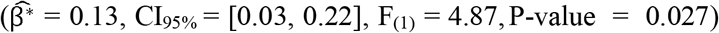. The significance of other variables remained unchanged. After backward selection, the model included net fishing selection, riverine net fishing licenses 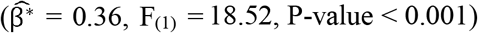, capelin 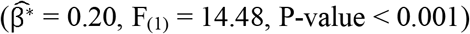, herring 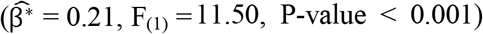 and krill 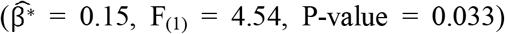 log-biomass. The dispersion parameter was 0.88.

The quadratic trend in the net fishing selection was removed to refit the *vgll3* quasi-binomial model on the residuals 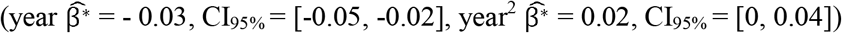. Riverine net fishing licenses, the net fishing selection and capelin log-biomass remained significant in the de-trended model (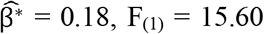 and P-value < 0.001, 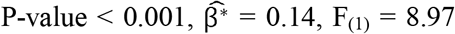 and 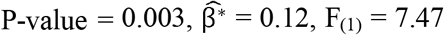 and P-value = 0.006; respectively) including a year effect (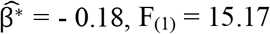 and P-value < 0.001).

